# Root-level loss of immunoglobulin and B-cell immune genes in clingfishes

**DOI:** 10.64898/2026.05.16.725622

**Authors:** Francisco Gambón Deza

## Abstract

Immunoglobulin genes are a central component of jawed-vertebrate adaptive immunity. A previous study showed that the blunt-snouted clingfish *Gouania willdenowi* lacks immunoglobulin genes and T-cell receptor gamma/delta loci, while retaining T-cell receptor alpha/beta genes, MHC genes, and *RAG1* /*RAG2*. Here I extend that observation to the family Gobiesocidae using all seven chromosome-level Gobiesocidae genome assemblies currently available. Manual tblastn and synteny-guided searches found no convincing immunoglobulin heavy-chain or light-chain loci in *G. willdenowi*, *Gouania pigra*, *Gobiesox punctulatus*, *Apletodon dentatus*, *Lepadogaster candolii*, *Lepadogaster purpurea*, or *Diplecogaster bimaculata*. Thus, the absence of antibody genes is best interpreted as a root-level character of clingfishes. The latest seven-species screen of 40 additional immune-associated genes shifts the broader interpretation in the same direction: the B-cell/adaptive core genes *CD79A*, *CD79B*, *CIITA*, *TNFRSF13B*, and *TNFSF13B* lack strong tblastn support in all sampled Gobiesocidae, and 37 of the 40 tested targets show an all-zero binary pattern at the presence threshold. Only *IL21R.1*, *TYROBP*, and *TNFRSF11A* show strong hits in one or more species. I therefore interpret the principal immune-gene erosion as occurring at or near the Gobiesocidae root rather than as a recent *Gouania*-specific process, while keeping weak, paralog-sensitive, and patchy loci provisional. RAG2 comparisons show a shared Gobiesocidae PHD-domain C-to-S replacement in the zinc-binding motif, with apparently intact *RAG2* coding sequence. A family-wide TRG/TRD screen did not recover *TRGV* V segments or accepted *TRDC* constant-region exons, but it did detect TRGC-like constant exons in several genomes. These TRGC-like sequences are probably not canonical TRG constant exons without further validation, so I treat the gamma/delta system as eroded or rearranged rather than as a complete root-level loss equivalent to the Ig loss. The RAG2 variant provides a plausible molecular context for antigen-receptor remodeling, but it is not evidence that RAG genes are pseudogenized, because TCR alpha/beta, MHC genes, and *RAG1* /*RAG2* are retained. Gobiesocidae are therefore best described as a vertebrate family with ancestral loss of canonical immunoglobulin genes and associated root-level erosion of B-cell and immune-related genes, not as a lineage lacking adaptive immunity in its entirety.

**Highlights:**

- Seven chromosome-level Gobiesocidae genomes lack convincing canonical IgH and IgL loci.
- The strongest non-Ig losses map to the B-cell/adaptive core: *CD79A*, *CD79B*, *CIITA*, *TNFRSF13B*, and *TNFSF13B*.
- TCR alpha/beta, MHC genes, and *RAG1* /*RAG2* are retained, so Gobiesocidae should not be described as lacking adaptive immunity in full.
- A shared Gobiesocidae RAG2 PHD-domain C-to-S variant provides candidate molecular context for antigen-receptor remodeling.

## 1. Introduction

The adaptive immune system of jawed vertebrates is based on somatically diversified antigen receptors, the recombination machinery encoded by *RAG1* and *RAG2*, and antigen-presentation pathways centered on the major histocompatibility complex (MHC) [1–3]. In most vertebrates these components are treated as an interdependent and deeply conserved system. Teleost fishes, however, show extensive lineage variation in immune-gene repertoires, including expansions, contractions, and occasional losses of particular immune pathways [4–6].

*Gouania willdenowi*, a small Mediterranean clingfish of the family Gobiesocidae, was previously reported to lack immunoglobulin genes altogether [7]. Gobiesocidae taxonomy and phylogeny have been revised repeatedly, including recent molecular analyses of clingfish and *Gouania* diversification [8–12]. That study also reported absence of T-cell receptor gamma/delta loci, but not a total loss of T-cell receptors: TCR alpha/beta sequences were recovered. Similarly, MHC genes were detected, and *RAG1* /*RAG2* were present. The original observation should therefore be understood as a loss of antibody genes plus selected antigen-receptor changes, rather than as complete elimination of adaptive immunity.

The availability of additional chromosome-scale Gobiesocidae genomes now makes it possible to ask whether these losses are restricted to *G. willdenowi*, shared only within *Gouania*, or rooted deeper in clingfish evolution. This distinction is important. A species-specific loss would imply a recent and possibly idiosyncratic event; a family-wide pattern places the transition at or near the root of Gobiesocidae and changes the biological interpretation from terminal degeneration to an ancestral immune architecture.

Here I revisit this question using a conservative framework. First, I perform manual immunoglobulin-locus searches in all seven available Gobiesocidae genomes. Second, I use the latest seven-species Gobiesocidae tblastn screen to test whether immune and immune-associated gene losses detected from the *Salarias fasciatus*-*Gouania* comparison are instead shared across the family. Third, I keep TCR alpha/beta, TCR gamma/delta, MHC, and RAG as separate evidence classes so that antibody loss can be related to possible antigen-receptor remodeling without collapsing all adaptive immune components into a single absence claim. Blenniiform outgroups provide a suitable comparative frame for Gobiesocidae-scale inference [13, 14]. The goal is to define the evolutionary level at which the main losses occurred and to remove inconsistencies introduced by mixing root-level, terminal, and ambiguous calls.

## 2. Results

### 2.1. Immunoglobulin genes are absent from all examined Gobiesocidae

The prior study established the absence of immunoglobulin genes in *G. willdenowi* by manual searches and synteny analysis [7]. I extended this approach to six additional Gobiesocidae genomes: *Gouania pigra*, *Gobiesox punctulatus*, *Apletodon dentatus*, *Lepadogaster candolii*, *Lepadogaster purpurea*, and *Diplecogaster bimaculata*. These assemblies, together with *G. willdenowi*, represent the currently available chromosome-scale genomic sampling of the family.

Manual searches targeted immunoglobulin heavy-chain constant regions, heavy- and light-chain V segments, and genomic regions syntenic to Ig loci in related teleosts. Searches used tblastn against raw genomic sequence, inspection of candidate hits, and comparison of flanking genes. No species yielded a convincing coding IgH, IgK, IgL, or teleost-specific light-chain locus. I therefore interpret the absence of canonical functional immunoglobulin loci as a root-level character of Gobiesocidae (Fig. 5). This moves the main evolutionary event from a terminal *Gouania* change to a basal transition before the diversification of the sampled clingfish lineages.

This result should not be read as a claim that every molecular remnant of the ancestral Ig loci has been eliminated. A systematic pseudogene screen, designed to recover highly eroded fragments with reduced identity, disrupted reading frames, or premature stop codons, was not performed across all seven genomes. The present inference is therefore a functional-locus inference: canonical coding IgH/IgL loci are not recovered, whereas deeply eroded relics remain a testable possibility. Such relics, if found within expected syntenic intervals, would strengthen the interpretation of evolutionary loss rather than assembly absence. This distinction is especially relevant because the TRG/TRD screen detects TRGC-like subthreshold exons in some genomes, consistent with the possibility that parts of the antigen-receptor region may persist as eroding fragments.

This conclusion is deliberately limited to immunoglobulin genes. It should not be extended to all antigen-receptor loci. In *G. willdenowi*, the previous study detected TCR alpha/beta sequences, while TCR gamma/delta were not detected [7]. The available data therefore support root-level loss of antibody genes across Gobiesocidae and absence of TCR gamma/delta in *G. willdenowi*, but they do not support a blanket statement that all TCR loci are absent from all Gobiesocidae.

I therefore keep the two TCR arms as separate evidence classes. To extend the V-exon evidence across Gobiesocidae, I applied Vgeneextractor to the seven genome assemblies and classified the final V candidates by neural-network assignment. CHfinder was applied in parallel to search for immunoglobulin heavy-chain constant-region exons. The Vgeneextractor screen recovered TRA/TRD-like and *TRBV* candidates in all sampled species, with *NITR* candidates recovered in the newly screened genomes where that class was scored, but it did not recover accepted light-chain V segments or *TRGV* candidates (Table 1). The two *IGHV* -like candidates found in *L. purpurea* are located on the same genomic segment as the *TRBV* cluster, and are therefore not interpreted as evidence for an immunoglobulin heavy-chain locus. CHfinder did not recover accepted immunoglobulin M, D, or T heavy-chain constant exons in any of the seven genomes.

**Table 1:**
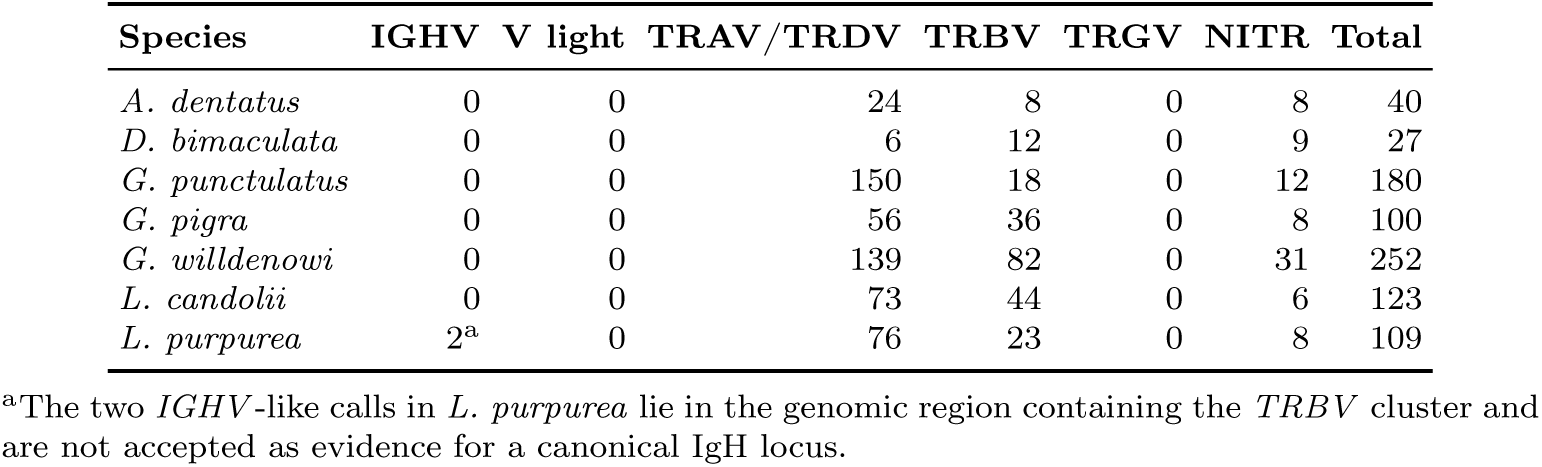
Vgeneextractor V-class calls across the seven Gobiesocidae genomes. Counts are based on the final selected V-exon candidates, with the *G. willdenowi* row using the values reported in the previous article for that species. Because Vgeneextractor does not distinguish *TRAV* from *TRDV*, the *TRAV* /*TRDV* column is interpreted as a combined alpha/delta V-like class.

| Species | IGHV | V light | TRAV/TRDV | TRBV | TRGV | NITR | Total |
| --- | --- | --- | --- | --- | --- | --- | --- |
| *A. dentatus* | 0 | 0 | 24 | 8 | 0 | 8 | 40 |
| *D. bimaculata* | 0 | 0 | 6 | 12 | 0 | 9 | 27 |
| *G. punctulatus* | 0 | 0 | 150 | 18 | 0 | 12 | 180 |
| *G. pigra* | 0 | 0 | 56 | 36 | 0 | 8 | 100 |
| *G. willdenowi* | 0 | 0 | 139 | 82 | 0 | 31 | 252 |
| *L. candolii* | 0 | 0 | 73 | 44 | 0 | 6 | 123 |
| *L. purpurea* | 2<sup>a</sup> | 0 | 76 | 23 | 0 | 8 | 109 |

<sup>a</sup>The two *IGHV*-like calls in *L. purpurea* lie in the genomic region containing the *TRBV* cluster and are not accepted as evidence for a canonical IgH locus.

**Table 2:**
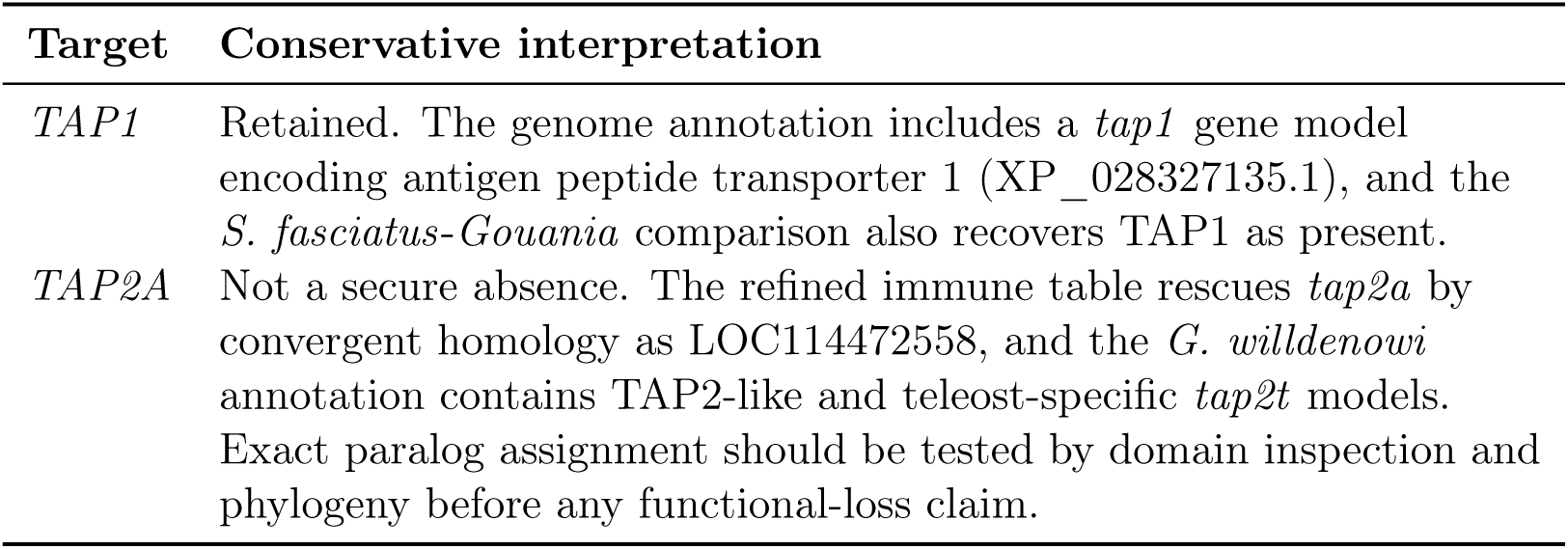
Clarification of TAP calls in *Gouania willdenowi*.

**Table 3:** Focused CD79 remnant and mechanism check in *Gouania willdenowi*.

| Locus | Gouania | Salarias | Mechanism |
| --- | --- | --- | --- |
| *CD79A* | 4,236 bp | 3,818 bp | No significant genome-wide or local tblastn remnant; adjacent conserved flanks with no intervening coding gene indicate focal deletion or complete local erosion. |
| *CD79B* | 19,829 bp | 3,739 bp | No significant genome-wide or local tblastn remnant; the interval between retained flanks contains two non-CD79 genes, indicating CD79B loss with local reorganization or insertion. |

**Table 4:** Conservative summary of refined candidate absences and the latest seven-species Gobiesocidae immune-gene screen.

| Category | Count |
| --- | --- |
| Candidates reanalysed by homology | 1,036 |
| Rescued by convergent homology | 844 |
| Rescued by a single reference | 16 |
| Weak convergent homology | 60 |
| Weak or ambiguous | 10 |
| High-confidence absences | 106 |
| Immune candidates reanalysed | 19 |
| Immune high-confidence absences | 5 |
| Additional immune-associated genes screened across Gobiesocidae | 40 |
| Gene-by-species tblastn calls | 280 |
| Strong present calls at the binary threshold | 10 |
| Targets with all-zero binary pattern across seven species | 37 |
| Root-compatible B-cell/adaptive core targets | 5 |

**Table 5:** *Salarias*-anchored immune-associated candidates screened across Gobiesocidae. These genes are present in *Salarias fasciatus* but lack a high-confidence DNA counterpart in *G. willdenowi* after protein and genome-level filtering. The latest seven-species tblastn screen places most of them as root-compatible absences, while weak and paralog-sensitive rows remain provisional.

| Functional group | Candidate genes and interpretation |
| --- | --- |
| B-cell / adaptive core | *CD79A*, *CD79B*, *CIITA*, *TNFRSF13B*, *TNFSF13B*.<br>Highest-confidence root-compatible immune losses; coherent erosion of BCR signaling, B-cell survival, and MHC class II regulation. |
| Cytokines and cytokine receptors | *IL13RA2*, *IL21R.1*, *IL1RL1*, *IL1RL2*, *CRFB1*, *CRFB2*, *IL22*, *IL17RB*, *IL17RC*, *IL12RB2L*, *CSF2RB*, plus *IFNGR1*, *IFNGR2*, and *IL23R* as orthology-sensitive candidates. Mostly root-compatible absence at the binary threshold; receptor-family calls require paralogy-aware validation. |
| Signaling and inflammatory mediators | *TIRAP*, *IRF1B*, *IRF2B*, *TYROBP*, *C4B*, *ALOX5AP*, *LTC4S*.<br>Candidate root-level losses affecting Toll/IL-1 adaptor signaling, interferon regulatory factors, complement, and eicosanoid metabolism; *TYROBP* has strong hits in two species and is therefore not a clean root-level call. |
| Other immune-associated genes | *IL16*, *CKLF*, *PRC1B*, *C2CD2L*, *SPAG7*, *LCP2A*, *MEA1*, *CD164*, *IGLDCP*, *ILF2*, *CD68*, *CD44B*, *TNFRSF9A*, *TNFRSF11A*. Broader immune-associated or immune-keyword candidates; useful for pathway context but not all are core immune genes; *TNFRSF11A* is present in six species and is not part of the root-level loss set. |

This software screen supports the manual interpretation: TCR alpha/beta-like V-exon signals are retained, whereas canonical IgH/IgL loci remain absent. The Vgeneextractor classifier does not distinguish *TRAV* from *TRDV* ; consequently, the *TRAV* /*TRDV* column in Table 1 is treated as a combined alpha/delta V-like category and not as a resolved *TRAV* -only call. In the prior *G. willdenowi* study, *TRAC* and *TRBC* were reported as present (Supplementary Fig. S3), whereas *TRGC* and *TRDC* were not detected in the genome/transcriptome searches [7]. This supports retention of TCR alpha/beta in *G. willdenowi* and non-detection of TCR gamma/delta in that species.

I then performed the required family-wide TRG/TRD constant-region screen with 155 TRGC and 167 TRDC protein queries, tblastn against each Gobiesocidae assembly, candidate exon extraction, and neural-network classification. The *S. fasciatus* outgroup control was strongly positive for TRGC, showing that the screen can recover TRG constant regions when they are present (Table S1). Specifically, *S. fasciatus* yielded 3,270 TRGC tblastn hits, 325 accepted TRGC candidates, and a best direct hit with 100% identity and 100% query coverage. No Gobiesocidae genome produced a strict strong-direct TRGC or TRDC tblastn call, and no accepted TRDC constant-region exon was recovered. However, TRGC-like constant exons were recovered in *A. dentatus*, *L. candolii*, and *L. purpurea*, with homologous subthreshold TRGC-like exons in both *Gouania* genomes. A classifier-positive *D. bimaculata* candidate was low-complexity and repeat-like, and was not treated as robust TRGC evidence. Thus the family-wide search resolves the root question conservatively: the Ig loss can be assigned to the Gobiesocidae root with high confidence, but complete TCR gamma/delta loss cannot. The current evidence supports loss or non-detection of *TRGV* and *TRDC*, together with persistence of TRGC-like remnants or exons in several lineages. These TRGC-like sequences should probably not be interpreted as canonical TRG constant exons at present: their relationship to a functional gamma/delta TCR locus remains unresolved and requires targeted locus-level and transcript-level validation.

### 2.2. Gouania willdenowi retains RAG, MHC and TCR alpha/beta signals

Because immunoglobulin genes are absent, genes involved in V(D)J recombination and antigen presentation require careful interpretation. *RAG1* and *RAG2* are present in the current *G. willdenowi* annotation and should not be treated as pseudogenes without explicit functional evidence. Likewise, MHC genes are detectable in *G. willdenowi*, and the presence of TCR alpha/beta sequences indicates that the cellular arm of adaptive immunity has not simply disappeared. These observations constrain the evolutionary interpretation: Gobiesocidae are not best described as lacking adaptive immunity in toto, but as lacking canonical immunoglobulin genes and showing root-level erosion of B-cell and immune-associated genes, with selected antigen-receptor components remodeled rather than uniformly deleted.

A separate MHC class II pathway screen refined this point. MHC class II alpha and beta structural genes were recovered in all seven Gobiesocidae genomes, as were *CTSS* and the basal class II transcription-factor components *RFX5*, *RFXANK*, *RFXAP*, *NFYA*, *NFYB*, and *NFYC* ; *CD74* was recovered in six of seven species. The consistent exception was *CIITA*: no sampled Gobiesocidae genome produced a strong intact *CIITA* hit, with only weak or fragmentary support in most species and no above-threshold support in *D. bimaculata*. Thus, the class II result is not a loss of the MHC class II structural genes, but a family-wide absence or severe degradation of the dedicated *CIITA*-dependent transcriptional regulator of canonical MHC class II expression [15].

### 2.3. A shared RAG2 PHD-domain variant accompanies antibody-gene absence

To add a protein-level check of the recombination machinery, I compared the RAG2 PHD zinc-binding motif across the seven Gobiesocidae genomes and eight blenniiform outgroups. *RAG2* was recovered as an apparently intact coding sequence in the sampled Gobiesocidae, without internal stop codons in the recovered open reading frames. The RAG2 PHD finger is functionally important because it couples chromatin recognition to V(D)J recombination, especially through H3K4me3 binding [16–18]. The outgroups retain the canonical GYWIKCC motif at the core of the PHD zinc-binding region, whereas all seven Gobiesocidae share a C-to-S replacement at the second cysteine of this pair, producing a GYWIKCS motif (Fig. 6). In the available sample, the CC motif co-occurs with Ig-positive outgroups and the CS motif co-occurs with Ig-negative Gobiesocidae.

This pattern is useful because it is a shared protein-level character of the Gobiesocidae antibody-loss background. It also provides the most direct molecular bridge between antibody loss and the mixed TCR gamma/delta pattern: if the PHD-domain replacement altered RAG2 chromatin coupling or recombination efficiency, Ig loci and TRG/TRD loci could have experienced different selective or functional constraints from retained TCR alpha/beta loci. This remains a hypothesis, not a demonstrated mechanism. The conservative interpretation is that Gobiesocidae retain RAG-mediated recombination machinery, while RAG2 carries a root-level PHD-domain variant that may reflect relaxed or altered selection after immunoglobulin-gene loss.

### 2.4. Seven-species immune-gene screen places B-cell erosion at the root

Figure 1 summarizes why the broad candidate-loss screen was not used directly as a final loss catalogue. An initial symbol-based comparison can overestimate gene loss, especially in teleosts where paralogy, lineage-specific naming, and annotation differences are common [19]. I therefore used the refined homology table gouania_missing_candidates_refined_by_homology.tsv to reassess candidate absences in *G. willdenowi*. Of 1,036 candidate genes in that table, 844 were rescued by convergent homology, 16 were rescued by a single reference, 60 showed weak convergent homology, 10 were weak or ambiguous, and 106 remained classified as high-confidence absences.

**Fig. 1.**
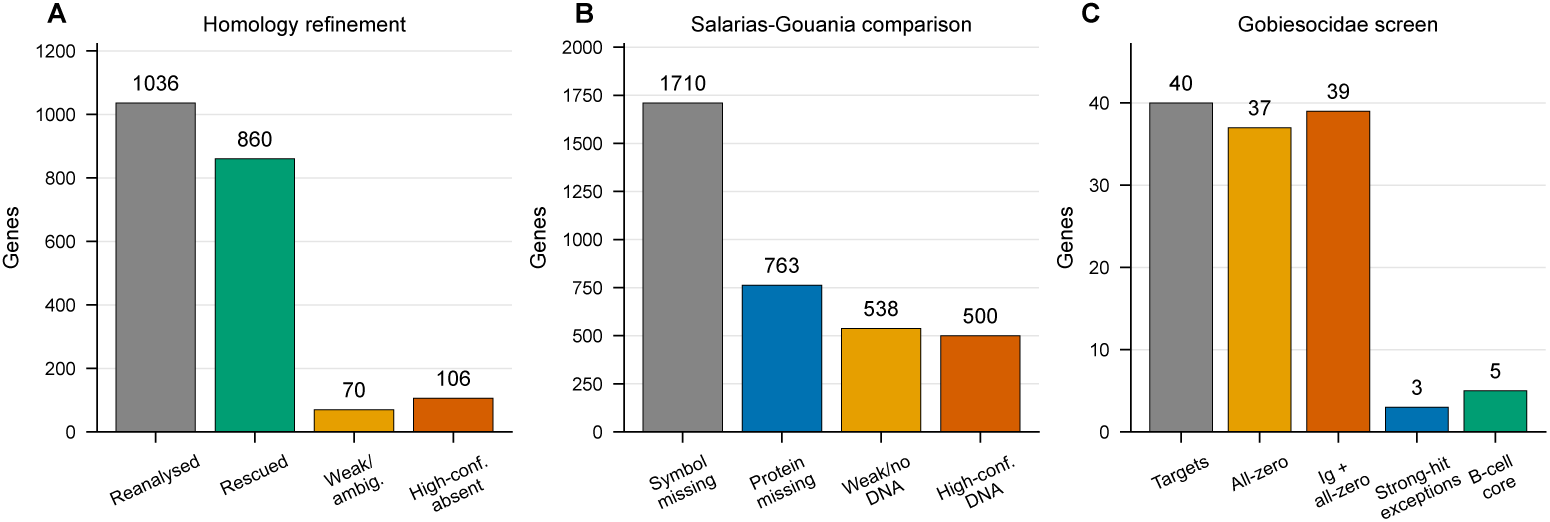
Conservative summary after root-level reassessment. Quantitative summary of candidate absence refinement, the *Salarias*-*Gouania* genomic comparison, and the latest Gobiesocidae root-level screen. The final panel shows that 37 of 40 additional immune-associated targets have all-zero binary calls across the seven sampled Gobiesocidae, producing 39 root-compatible targets when IgH and IgL are added. Only three screened rows have strong hits in one or more Gobiesocidae species, and the five B-cell/adaptive core genes form the cleanest non-Ig root-level set.

The latest Gobiesocidae analysis then tested whether immune and immune-associated candidates from this *Salarias*-anchored comparison were restricted to *G. willdenowi* or shared across the family. A targeted tblastn screen of 40 genes across the seven chromosome-level Gobiesocidae assemblies produced 280 gene-by-species calls: 10 strong present hits, 96 hits below the absence threshold, 72 weak hits below the presence threshold, and 102 cases with no tblastn hit. Weak hits were retained in the evidence table but encoded as zero in the binary matrix (Fig. 2). Thus, at the predefined presence threshold, 37 of the 40 tested targets show an all-zero pattern across the family; the only rows with strong hits in any species are *IL21R.1*, *TYROBP*, and *TNFRSF11A*.

**Fig. 2.**
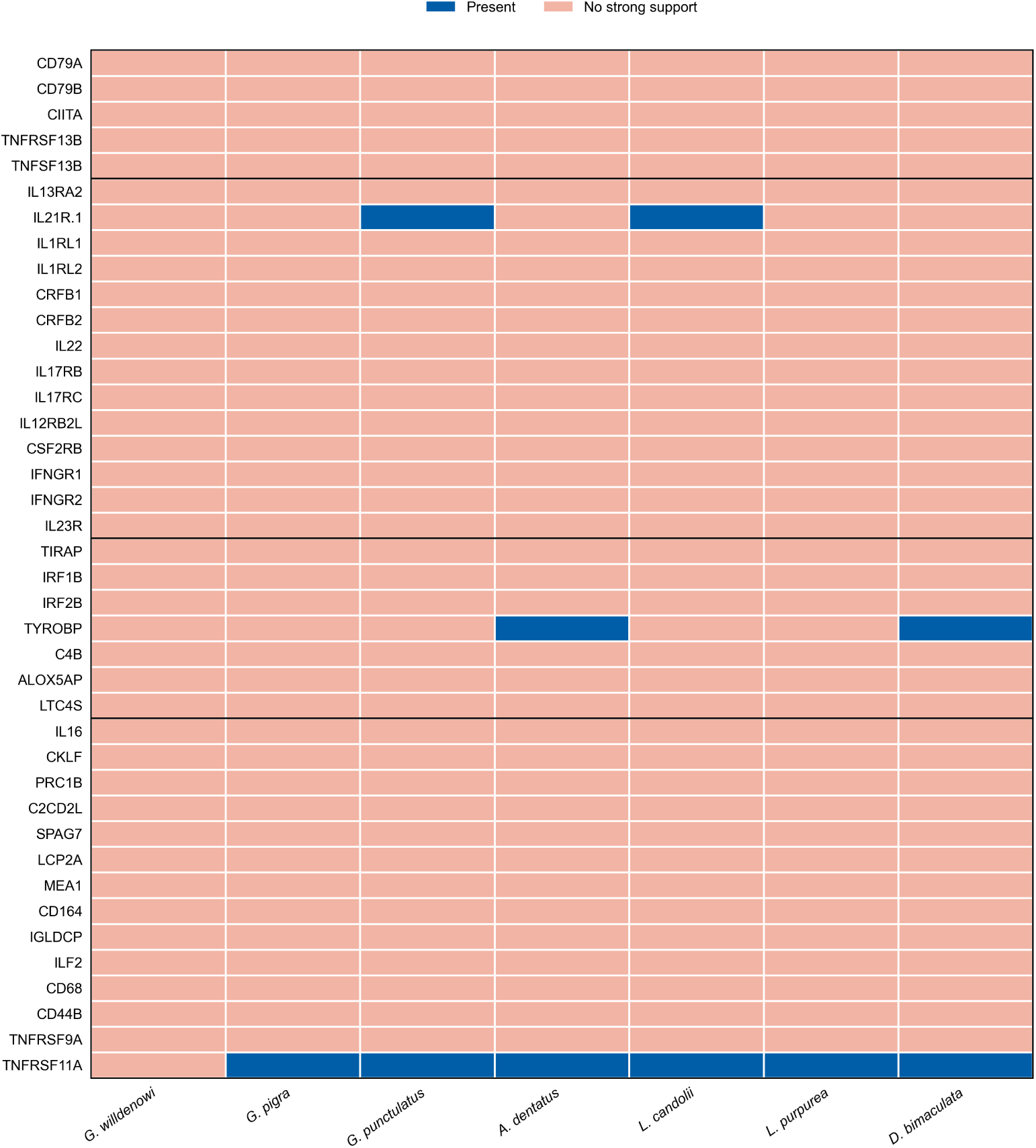
Gobiesocidae binary immune-gene matrix. Binary tblastn matrix for 40 immune-associated targets screened across the seven chromosome-level Gobiesocidae assemblies. Blue cells indicate strong presence calls meeting the predefined threshold; red cells indicate no strong support at that threshold. Weak subthreshold hits are retained in the evidence table but encoded as zero here. The dominant pattern is root-compatible absence, with strong-hit exceptions for *IL21R.1*, *TYROBP*, and *TNFRSF11A*.

The most important change is phylogenetic. Among the immune candidates that remain high-confidence absences in *G. willdenowi*, the B-cell/adaptive core genes *CD79A*, *CD79B*, *CIITA*, *TNFRSF13B*, and *TNFSF13B* now lack strong tblastn support in all sampled Gobiesocidae. These are therefore best interpreted as losses at or near the root of the family, not as a recent *Gouania*-only erosion. Functionally, they define an ancestral disruption of B-cell receptor signaling/survival and MHC class II regulation that is coherent with the family-wide loss of antibody genes [15, 20–22]. Broader all-zero cytokine, signaling, and immune-associated rows support additional root-level immune remodeling, but weak, rescued, or paralog-sensitive loci should still be treated as candidates until validated locus by locus.

The *TAP1* /*TAP2A* calls required separate inspection because an absence of either canonical TAP subunit would have a direct interpretation for MHC class I peptide loading [23, 24]. The symbol-level matrix scored both genes as absent in *G. willdenowi*, but this is not supported after annotation and homology review. The current *G. willdenowi* annotation contains *tap1* (XP_028327135.1), a TAP2-like model assigned to LOC114472558 (XP_028317681.1), a teleost-specific *tap2t* model (XP_028331989.1), and an additional low-quality TAP2-like model (XP_028298627.1). In the refined immune table, *tap2a* is rescued by convergent homology as LOC114472558, with approximately 72% identity across 97–99% of the compared references. I therefore treat *TAP1* as retained and *TAP2A* as rescued or paralog-ambiguous, not as a high-confidence absence.

Because *CD79A* and *CD79B* are central to B-cell receptor signaling, I also inspected their local genomic context. In the positive reference *S. fasciatus*, *CD79A* lies between conserved flanking genes including *ARHGEF1* and *LIPE* ; in *G. willdenowi*, the corresponding flanks are retained on chromosome 16 but the *CD79A* coding sequence is not detected in the intervening region (Supplementary Fig. S1). Similarly, the *CD79B* region is bracketed by conserved flanks including *SCN4A* and the downstream *PLEKHM1* side of the locus in the current *Salarias*-anchored comparison; these flanks are retained on *G. willdenowi* chromosome 19 while *CD79B* is not detected (Supplementary Fig. S2). Thus, for *G. willdenowi*, the *CD79A* and *CD79B* absence calls are supported by local synteny, not only by symbol-level annotation. The broader Gobiesocidae matrix scores both genes as absent in all seven sampled species, but locus-level flanking validation is currently strongest for the *Salarias*-*Gouania* comparison.

I then used the *S. fasciatus* CD79A and CD79B proteins as queries in an explicit remnant search against the full *G. willdenowi* genome and against the corresponding syntenic intervals. No hit met even a permissive remnant threshold (tblastn e-value ≤ 10^−3^ and query coverage ≥ 30%). The best genome-wide matches were short and nonsignificant (best e-values 0.45 for CD79A and 0.089 for CD79B), and the best local interval matches were weaker still. The local geometry therefore gives a more informative mechanism than the sequence remnants themselves. For CD79A, the retained *LIPEB* and *ARHGEF1* flanks are separated by only 4,236 bp in *Gouania*, comparable to the 3,818-bp *S. fasciatus* CD79A span, and no intervening protein-coding gene is annotated. This is most consistent with a small focal deletion or complete erosion of the CD79A coding interval rather than a retained pseudogene accumulating point mutations. For CD79B, the retained *SCN4AA* and *PLEKHM1* flanks are separated by 19,829 bp and contain two annotated non-CD79 genes in *Gouania*; this supports loss of the CD79B coding sequence with local reorganization or insertion, rather than a simple intact pseudogene. Thus, for these two BCR genes, the evidence favors physical removal or local locus restructuring over gradual mutation accumulation in a recognizable remnant.

The same logic was applied to the immunoglobulin heavy-chain locus. In *S. fasciatus*, the annotated IgH-rich region lies on chromosome 4 between the *CABP5* -like side of the locus and the downstream *NACC1A*/*IER2A*/*CACNA1AA* block, matching the microsynteny described for Ovalentaria IgH loci in the previous *Gouania* study. In *G. willdenowi*, these flanking genes are all retained on chromosome 8 (NC_041051.1), but the expected *NACC1A*-to-*CABP5* -like interval is only 155,994 bp, whereas the corresponding *S. fasciatus CABP5* -like-to-*NACC1A* span is about 1.00 Mb and contains the IgH cluster. The *Gouania* interval contains several non-IgH genes, but no canonical heavy-chain constant-region, heavy-chain V-segment, or IgH-cluster annotation. A focused tblastn search using the annotated *S. fasciatus* IgH constant-region proteins found no significant hit in the expected *Gouania* chromosome 8 interval; the best local match had e-value 0.062 and was therefore consistent with background Ig-domain similarity. Genome-wide permissive searches did recover weak matches outside the expected locus, mainly on chromosomes 18 and 22, where the annotation contains Ig-light-chain-like or TCR/Ig-domain-like features. These off-locus hits do not reconstruct a syntenic IgH locus. Thus, the IgH case supports a larger physical contraction/deletion of the antibody locus with local rearrangement, rather than retention of a recognizable IgH pseudogene at the ancestral chromosomal position.

### 2.5. The Salarias comparison identifies candidates whose phylogenetic level is tested across Gobiesocidae

A direct comparison between *G. willdenowi* and *Salarias fasciatus* provides an independent genomic layer. In the final comparison, *S. fasciatus* contains 12,552 named genes. Of these, 11,789 have protein-level support in *Gouania*, leaving 763 protein-missing candidates. After genome-level filtering, 538 candidates remain in weakly aligned or unaligned regions, and 500 are classified as high-confidence genes in DNA missing from *Gouania*.

These 500 genes are not randomly distributed. They include one large block, 51 small blocks, and 381 single-gene losses, with significant chromosomal heterogeneity (chi-squared *p* = 8.0 × 10^−9^). A permutation test also supports non-random clustering of large blocks (*p* ≃ 0.004). Figure 3 shows that the missing genes are concentrated in discrete chromosomal intervals rather than scattered uniformly across the genome, and Fig. 4 shows the same signal as local density peaks. These two views indicate that many candidate losses reflect structured genomic deletions or contractions, not random independent gene dropouts. Functionally, the high-confidence set is enriched for innate immune signaling, but most genes are outside canonical immune categories. This analysis supports a real genomic-loss signal in *G. willdenowi*, while reinforcing that the larger symbol-based list should be treated as a candidate screen rather than a final loss catalogue. In the revised interpretation, these *Salarias*-anchored candidates are not automatically considered terminal *Gouania* losses. Their evolutionary level is assigned only after the seven-species Gobiesocidae screen: shared absence across all sampled clingfishes is root-compatible, whereas strong hits in one or more species mark patchy or lineage-specific patterns.

**Fig. 3.**
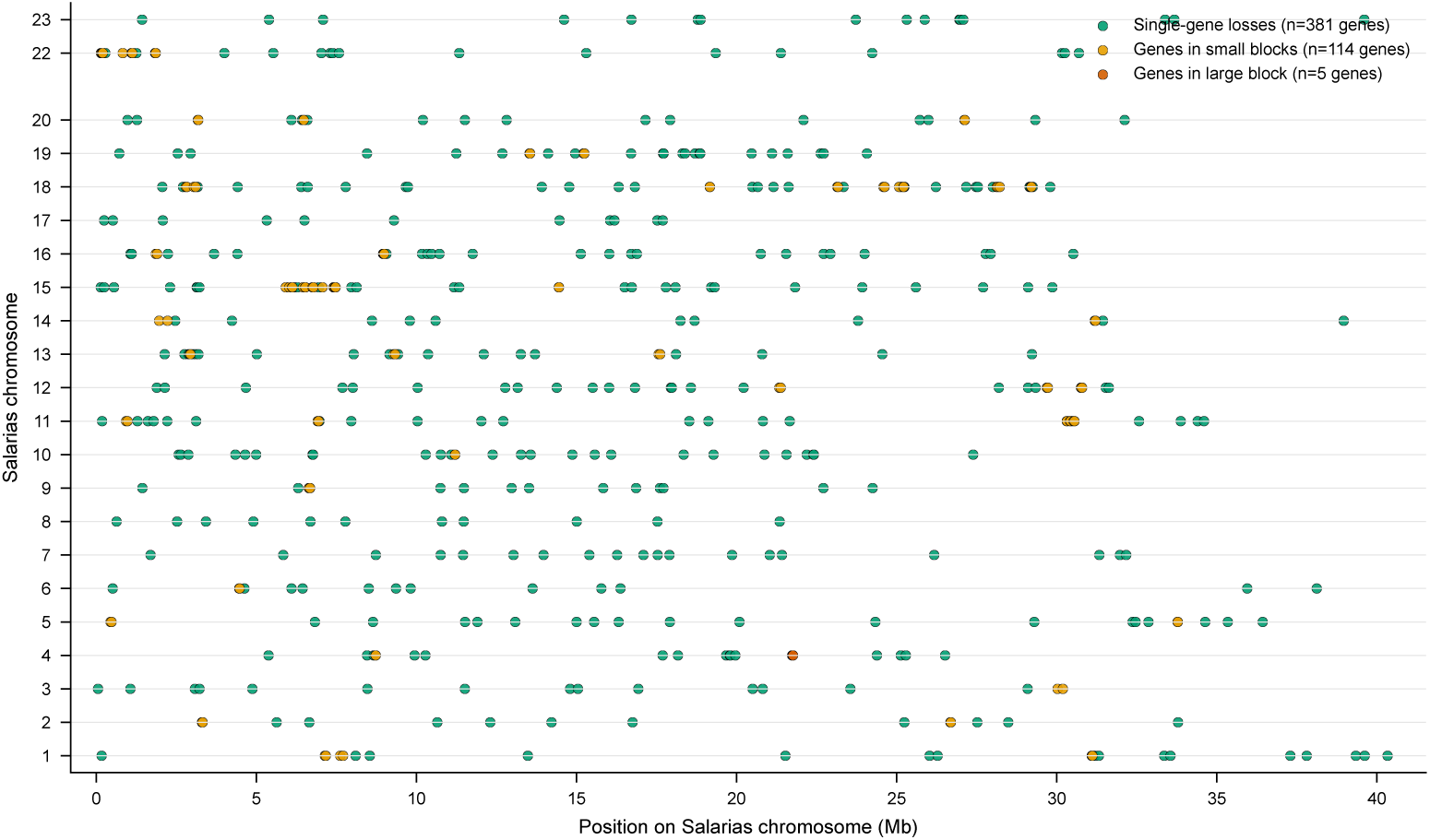
Chromosomal distribution of high-confidence *Gouania* missing-DNA genes. Positions of high-confidence genes from *S. fasciatus* that fall in regions with little detectable alignment to *G. willdenowi*. The figure summarizes genomic distribution rather than a final functional classification for every gene.

**Fig. 4.**
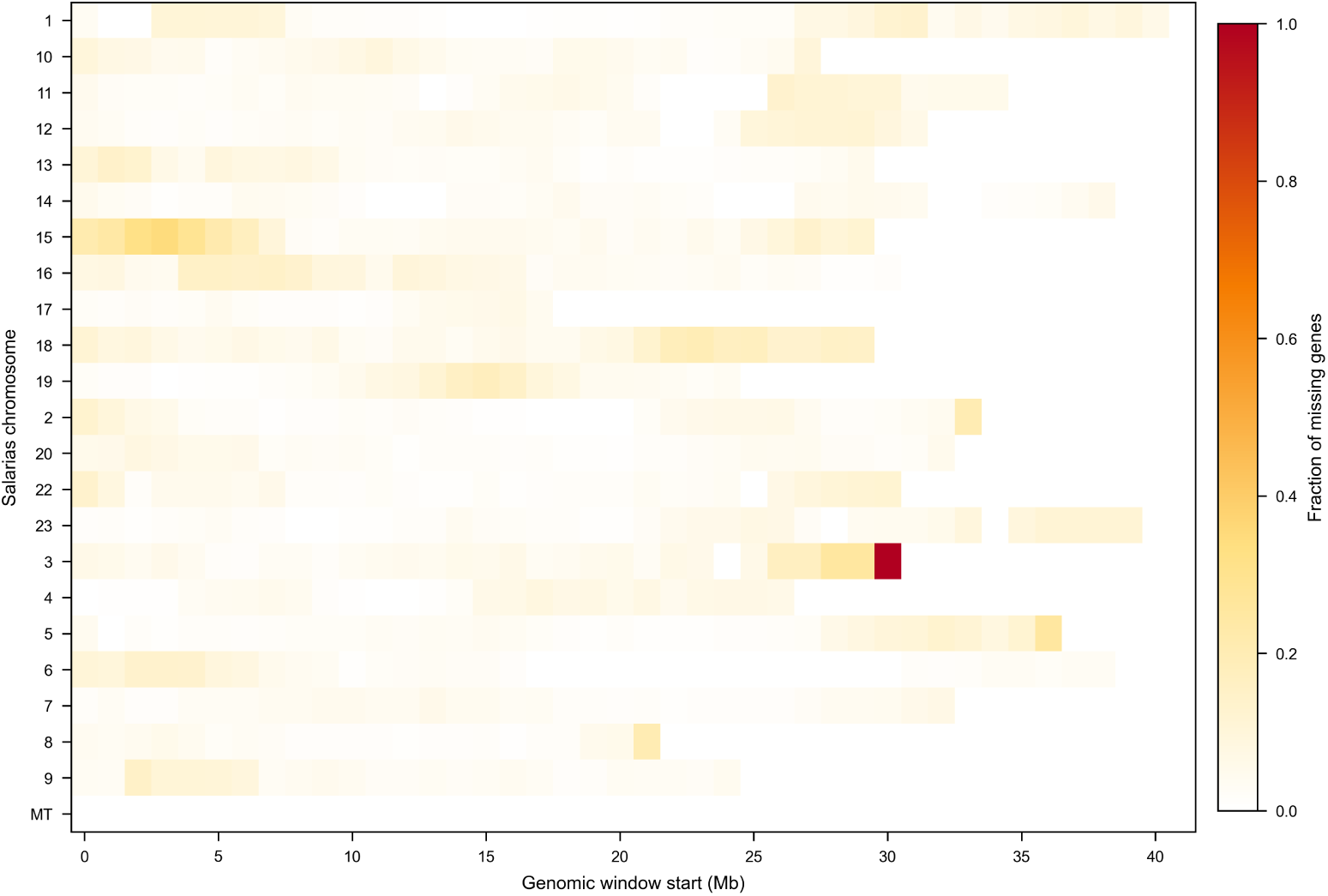
Density of candidate missing genes across chromosomes. Sliding-window density of high-confidence missing-DNA genes, showing that the signal is not explained by a single whole-chromosome loss.

The immune-related subset of this *Salarias*-anchored comparison contains 40 genes, which were subsequently screened across Gobiesocidae. These include cytokines and cytokine receptors (*IL13RA2*, *IL21R.1*, *IL1RL1*, *IL1RL2*, *CRFB1*, *CRFB2*, *IL22*, *IL17RB*, *IL17RC*, *IL12RB2L*, and *CSF2RB*), inflammatory and innate-signaling mediators (*TIRAP*, *IRF1B*, *IRF2B*, *TYROBP*, *C4B*, *ALOX5AP*, and *LTC4S*), and additional immune-associated genes such as *IL16*, *CKLF*, *LCP2A*, *CD68*, *CD44B*, *TNFRSF9A*, and *TNFRSF11A*. Thus, the direct genome comparison adds a biologically coherent layer of cytokine-receptor, signaling, and inflammatory candidates to the high-confidence B-cell and MHC-II regulatory set. The new family-wide screen indicates that most of these targets lack strong support in all sampled Gobiesocidae and are therefore compatible with root-level erosion. I still treat them as candidate losses, rather than validated functional absences, because several belong to paralogy-rich receptor families and some have weak subthreshold hits.

### 2.6. Most targeted immune-gene absences are root-compatible

The latest binary matrix changes the interpretation of the family-wide pattern (Fig. 2). Instead of a model dominated by recent *Gouania* or *G. willdenowi* losses, the targeted Gobiesocidae screen is dominated by shared non-detection. Thirty-seven of the 40 additional immune-associated targets lack strong tblastn support in all seven sampled species. Together with the manually verified absence of IgH and IgL loci, this places the main immune-gene reduction at or near the root of Gobiesocidae.

The cleanest root-compatible non-Ig set is the B-cell/adaptive core: *CD79A*, *CD79B*, *CIITA*, *TNFRSF13B*, and *TNFSF13B*. These genes connect directly to the antibody phenotype: *CD79A*/*CD79B* encode B-cell receptor signaling components, *TNFSF13B* /*TNFRSF13B* belong to the BAFF survival axis, and *CIITA* regulates MHC class II transcription. Their shared absence is therefore not an unrelated add-on to Ig loss, but part of the same ancestral loss-of-humoral-immunity signal.

The matrix still contains exceptions. *IL21R.1* has strong hits in *G. punctulatus* and *L. candolii*, *TYROBP* has strong hits in *A. dentatus* and *D. bimaculata*, and *TNFRSF11A* has strong hits in all sampled species except *G. willdenowi*. These rows should not be forced into the root-loss set. They mark either retained paralogs, lineage-specific differences, or targets requiring further orthology validation. The corresponding phylogenetic summary is shown in Fig. 5.

**Fig. 5.**
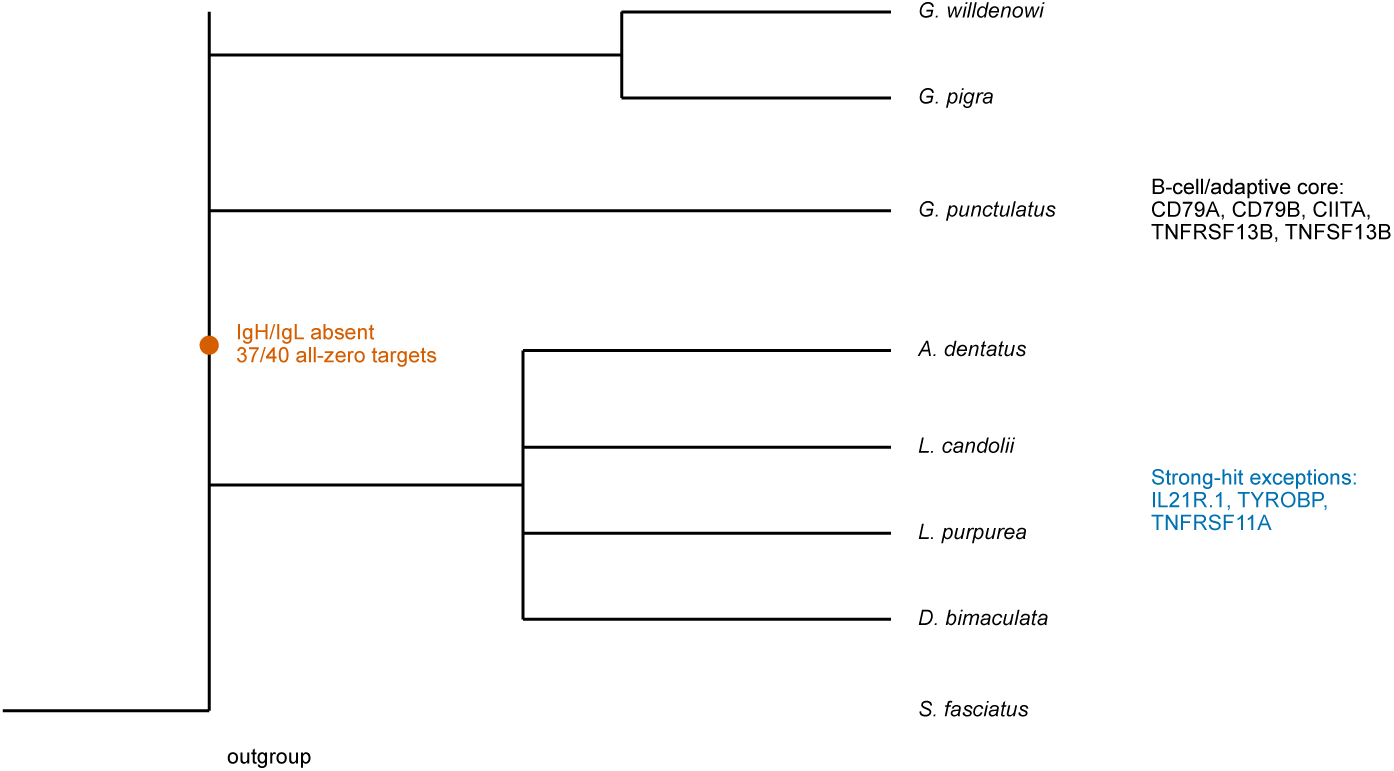
Root-level phylogenetic summary of immune-gene absence in Gobiesocidae. Phylogenetic representation of the revised interpretation. The principal event is placed at or near the root of Gobiesocidae: manual IgH/IgL absence plus 37 of 40 additional immune-associated targets with all-zero binary calls. The figure also lists the three screened rows with strong hits outside the root-loss set.

## 3. Discussion

### 3.1. A family of vertebrates without canonical immunoglobulin genes

The principal result of this study is that the immunoglobulin-gene absence first described in *G. willdenowi* extends to all examined Gobiesocidae and is most parsimoniously placed at or near the root of the family (Fig. 5). Because the conclusion is based on manual locus-level searches, not only on automated annotations, it is unlikely to be explained by poor gene prediction. Ig loci are difficult to annotate because they are repetitive and segmental, but the absence of convincing coding IgH or IgL candidates in seven chromosome-scale genomes is a strong signal.

The stronger evolutionary claim would be supported by a dedicated relic search around the expected IgH and IgL neighborhoods. The present data already argue against intact antibody loci, but they do not exclude short pseudogenic segments, isolated V-like fragments, or noncoding pieces of former constant regions. This is a limitation rather than a contradiction. If future analyses recover low-identity Ig remnants with frameshifts or stop codons in otherwise conserved flanking intervals, those remnants would favor an ancestral loss model over an assembly-gap model. Conversely, the complete absence of such remnants from multiple high-contiguity Gobiesocidae assemblies would indicate older or more complete erosion of the loci.

This finding makes Gobiesocidae an exceptional vertebrate family: the available genomes lack canonical antibody genes. The statement should be kept precise. The evidence does not require the disappearance of all adaptive immunity. *G. willdenowi* retains TCR alpha/beta signals, MHC genes, and RAG genes. The evolutionary problem is therefore more specific and more interesting: how can a vertebrate family persist without conventional immunoglobulin-mediated humoral immunity while retaining other components of adaptive and innate immune function?

### 3.2. RAG retention and mixed TCR gamma/delta evidence define a narrower model

The RAG data sharpen this conclusion. A shared RAG2 PHD-domain C-to-S replacement separates sampled Gobiesocidae from blenniiform outgroups, and it co-occurs with the absence of canonical Ig genes (Fig. 6). Nevertheless, *RAG1* and *RAG2* should be treated as retained unless functional experiments show otherwise. The RAG2 motif therefore provides a molecular signature of the Gobiesocidae antibody-loss background, but not evidence for wholesale loss of V(D)J recombination.

**Fig. 6.**
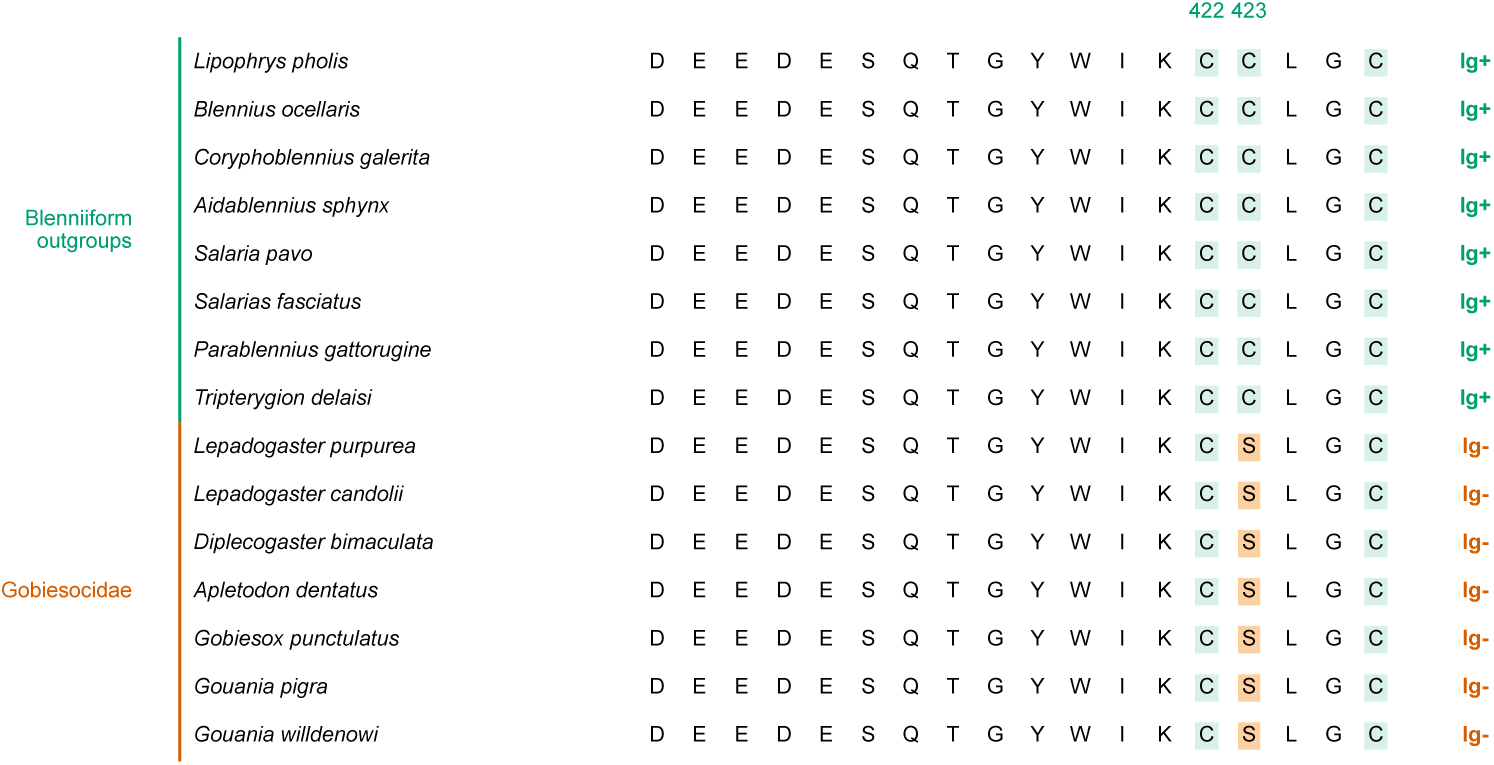
RAG2 PHD-domain motif in Gobiesocidae and blenniiform outgroups. Alignment of the RAG2 PHD zinc-binding motif surrounding positions 422–423, numbered by *Lipophrys pholis* RAG2. Blenniiform outgroups retain the GYWIKCC motif and have detectable immunoglobulin genes, whereas all sampled Gobiesocidae carry GYWIKCS and lack canonical Ig genes. The association is interpreted as a conserved RAG2 motif variant in the antibody-loss background, not as evidence that RAG2 is absent.

The TCR gamma/delta evidence is important but no longer supports a simple complete-loss statement. Vgeneextractor did not recover *TRGV* in any sampled Gobiesocidae genome, and the family-wide constant-region screen did not recover accepted *TRDC* exons. In contrast, TRGC-like constant exons persist in several genomes, including accepted classifier calls in *A. dentatus*, *L. candolii*, and *L. purpurea*, and homologous subthreshold candidates in *Gouania*. These sequences are best described as TRGC-like and pending study, because the present data do not show that they are canonical TRG constant exons embedded in an intact gamma/delta TCR locus. Antibody genes and TCR gamma/delta loci both depend on V(D)J recombination, whereas TCR alpha/beta signals are retained in *G. willdenowi*. The shared RAG2 PHD-domain replacement is therefore a plausible candidate modifier of the antigen-receptor landscape, but the direction of causality is unresolved. It could have preceded Ig loss, followed it under relaxed selection, or simply mark a lineage in which humoral immunity and parts of the gamma/delta receptor system were no longer maintained. The manuscript should state this as a mechanistic hypothesis, not as proof that the RAG mutation caused antibody loss.

### 3.3. The B-cell axis is the most defensible root-level target

Among non-Ig immune genes, the most robust root-compatible losses are *CD79A*, *CD79B*, *CIITA*, *TNFRSF13B*, and *TNFSF13B*. *CD79A* and *CD79B* encode the signaling components of the B-cell receptor complex, while *TNFSF13B* (BAFF) and *TNFRSF13B* are central to B-cell survival and maturation [20–22]. *CIITA* adds a class II antigen-presentation regulatory component [15]. Their shared absence is coherent with the loss of immunoglobulin genes and supports a basal dismantling of the canonical B-cell/humoral axis. This B-cell-centered result is much stronger than broader claims about full elimination of MHC, RAG or all TCR pathways.

The MHC class II finding is therefore regulatory rather than structural. The retention of MHC class II alpha/beta genes, *CTSS*, most *CD74* calls, and the basal RFX/NF-Y promoter machinery argues against describing Gobiesocidae as MHC-II-null. In contrast, the family-wide lack of an intact *CIITA* candidate suggests that canonical inducible MHC class II expression may be strongly reduced, lost, or replaced by a noncanonical regulatory route. Testing that possibility will require expression data from immune tissues or stimulated cells, together with locus-level validation of class II transcripts and peptide-presentation function.

### 3.4. Innate immune genes provide candidate compensatory capacity

The absence of antibody genes raises the question of whether other immune systems could buffer the loss of conventional humoral immunity. The *G. willdenowi* annotation does not show a genome devoid of immune defense. A simple protein-coding gene survey of the RefSeq feature table recovers multiple innate immune and antimicrobial categories: 15 toll-like-receptor entries, including *TLR1*, *TLR3*, *TLR5* /*TLR5B*, *TLR7*, *TLR8*, *TLR9*, *TLR13*, *TLR18*, and *TLR21* ; 95 NLR/NOD-like entries, including *NOD1*, *NOD2*, *NLRC5*, *NLRX1*, and many *NLRC3* -like models; 42 complement-related genes, including *C3* -like, *C5*, *C6*, *C7*, *C8*, *C9*, factor B, factor D, factor H, and factor I entries; 43 lectin-family entries; 58 interferon-related entries, including *STING1*, *IFIH1*, *RSAD2*, *ISG20*, Mx-like genes, and multiple interferon regulatory factors; and 20 antimicrobial-peptide or peptide-like entries, including *HAMP* /hepcidin-like genes, *LEAP2*, NK-lysin-like, and lysozyme genes. TRIM-family antiviral/innate candidates (*TRIM21*, *TRIM25*, *TRIM39*, and *TRIM47*) are also numerous in the annotation.

The classical complement pathway requires a recognizable C1 complex, including C1q A/B/C chains and the associated C1r/C1s proteases. In the refined *Salarias*-anchored homology table, the *c1qa*, *c1qb*, and *c1qc* candidates are not recovered as clean canonical C1q orthologues; their best matches point to C1q/TNF-like paralogues, and the calls remain weak or ambiguous. The *c1r* and *c1s.2* rows are rescued only through convergent homology to *masp1* -like sequences. Therefore, downstream complement capacity appears broadly retained, but the canonical antibody-triggered classical pathway is not demonstrated as conserved in *G. willdenowi* and should be treated as unresolved or potentially remodeled until the C1 locus is validated directly.

These retained genes are not proof that innate immunity functionally compensates for the loss of immunoglobulins. The survey is annotation-based, and several NLR and TRIM entries are paralog-rich, partial, or labelled “like”. Nevertheless, it provides a plausible biological context for survival without canonical antibodies: Gobiesocidae appear to retain broad pattern-recognition, complement, interferon, lectin, and antimicrobial-peptide toolkits. The defensible model is therefore not immune collapse, but a shift away from antibody-mediated humoral immunity toward retained and possibly remodeled innate and cellular immune defenses.

### 3.5. TAP calls do not support abolition of MHC class I presentation

The TAP1/TAP2 heterodimer normally imports proteasome-derived peptides into the endoplasmic reticulum for MHC class I loading [23, 24]. A true loss of either essential canonical subunit could strongly reduce conventional MHC class I surface presentation. The present data, however, do not support a TAP-null interpretation for *G. willdenowi*. *TAP1* is annotated, TAP2-like or *tap2t* models are present, *TAP2A* is rescued by convergent homology, and multiple MHC class I-like heavy-chain proteins are annotated in the same genome.

I therefore do not infer that class I antigen presentation is abolished in *G. willdenowi*. The defensible conclusion is narrower: the MHC class I processing pathway may be remodeled, and the exact identity of the TAP2-like component should be validated. Functional loss would require additional evidence, including expression of the TAP models, intact transporter-domain architecture, phylogenetic assignment of *TAP2A*/*tap2t* paralogs, and direct assays of peptide transport or surface MHC class I presentation.

### 3.6. Why the broader root-level catalogue must be conservative

The analyses show why a conservative interpretation is necessary. A large initial candidate list collapses substantially after homology rescue; many immune genes initially scored as absent are recoverable or ambiguous. A direct genome comparison with *Salarias* still supports hundreds of high-confidence missing genes, but these calls are not identical to the symbol-level list and should be interpreted as genomic absence candidates requiring targeted inspection for individual high-value loci.

The latest family-wide matrix nevertheless changes the direction of the argument. Most targeted immune-associated genes are not terminal *Gouania* losses in the binary screen; they are shared absences across the sampled family. The caution now concerns validation of individual loci, not the phylogenetic level of the dominant signal. Some rows are clean (*CD79A*, *CD79B*, *CIITA*, *TNFRSF13B*, *TNFSF13B*), whereas others are patchy or paralog-sensitive. Patchiness can reflect true retained paralogs, repeated loss, assembly gaps, divergent sequence, threshold effects, or annotation differences. The manuscript should therefore present broad non-Ig immune erosion as root-compatible and hypothesis-generating, while reserving strongest status for manually verified Ig loss and the B-cell/adaptive core. This caution is especially important for the broad tblastn matrix because its presence threshold was intentionally permissive. A 25% amino-acid identity cutoff with 40% query coverage can capture remote homologs and shared-domain matches in paralogy-rich immune families. I therefore use that matrix as a screen for root-compatible patterns, not as a stand-alone orthology test; weak or isolated positives require synteny, domain architecture, reciprocal homology, or manual locus inspection before they are interpreted as retained genes.

### 3.7. An evolutionary model

The most parsimonious model is that canonical immunoglobulin genes were lost at or near the base of Gobiesocidae, before the diversification of the sampled lineages. The same basal transition also involved loss or severe erosion of the B-cell/adaptive core, including *CD79A*, *CD79B*, *CIITA*, *TNFRSF13B*, and *TNFSF13B*. Many additional cytokine, signaling, and immune-associated targets follow the same all-zero binary pattern and are best treated as root-compatible candidate losses. A small number of rows, including *IL21R.1*, *TYROBP*, and *TNFRSF11A*, show strong hits in one or more species and should be kept outside the root-loss set unless future orthology work changes their status. The shared RAG2 PHD-domain variant and the mixed TCR gamma/delta evidence fit this ancestral remodeling model, but they should be interpreted as candidate mechanistic context rather than as primary proof of adaptive-immune collapse.

## 4. Materials and Methods

### 4.1. AI-assisted analysis and writing support

OpenAI Codex 5.4 and 5.5 were used under the supervision of the author to assist with code execution, file organization, LaTeX editing, and manuscript revision. All biological interpretations, methodological decisions, and final text were reviewed and approved by the author.

### 4.2. Genome assemblies

The study used the seven currently available chromosome-level Gobiesocidae genome assemblies: *Gouania willdenowi* (GCF_900634775.2), *Gouania pigra* (GCA_982345305.1), *Gobiesox punctulatus* (GCA_977017485.1), *Apletodon dentatus* (GCA_965542115.1), *Lepadogaster candolii* (GCA_965278445.2), *Lepadogaster purpurea* (GCA_965274325.2), and *Diplecogaster bimaculata* (GCA_976916555.1). The principal outgroup/comparison genome for direct gene-loss analysis was *Salarias fasciatus* (GCF_902148845.1). Species selection and outgroup placement followed the available Gobiesocidae and blenniiform phylogenetic framework [10–14].

### 4.3. Manual immunoglobulin searches

Immunoglobulin searches were performed manually using tblastn against raw genomic assemblies [25], protein queries from related teleosts, and inspection of syntenic regions surrounding known Ig loci in outgroup species. Candidate hits were evaluated for Ig-domain structure, genomic context, and consistency with heavy- or light-chain architecture. Absence calls were accepted only when no convincing coding region or syntenic Ig locus was recovered. These searches were designed to test for canonical coding Ig loci, not to exhaustively catalogue pseudogenes. A dedicated pseudogene/remnant analysis would require relaxed fragment-level searches in the expected flanks, translation of low-identity hits in all frames, and manual scoring of frameshifts, premature stop codons, splice disruption, RSS decay, and local synteny.

The manual Ig-locus assessment was complemented with Vgeneextractor and CHfinder. Vgeneextractor was used to recover and classify V-exon candidates from each Gobiesocidae genome into immunoglobulin, TCR, and NITR-like classes. Because this classifier does not separate *TRAV* from *TRDV*, TRA/TRD-like candidates were reported as a combined category. CHfinder was used to search for immunoglobulin heavy-chain constant exons corresponding to IgM, IgD, and IgT; isolated low-confidence candidates were not accepted unless they formed a coherent immunoglobulin constant-region signal.

To test whether TCR gamma/delta constant-region loss could be assigned to the family root, I added a separate TRG/TRD constant-region screen. Protein queries were assembled from local TRGC/TRDC reference sets, excluding records labelled as background or failed TRGC examples, resulting in 155 nonredundant TRGC and 167 nonredundant TRDC queries. These queries were searched against each Gobiesocidae assembly with tblastn. The initial tblastn pass was deliberately inclusive and was used to define candidate windows, not to assign presence by itself. Candidate splice-bounded exons surrounding hits were translated and classified with the same neural-network constant/V classifier. A candidate was accepted automatically only when it was labelled *TRGC*, *C_gamma*, or *TRDC* with probability at least 0.5, and strict direct tblastn evidence required e-value ≤ 10^−5^, amino-acid identity ≥ 30%, query coverage ≥ 50%, and bit score ≥ 45. The *S. fasciatus* genome was included as an outgroup control, and the complete control-and-family summary is provided in Table S1 and source file table_s1_trg_trd_screen.tsv.

### 4.4. TCR alpha/beta and gamma/delta evidence handling

TCR evidence was not pooled into a single all-TCR absence category. I treated the prior *G. willdenowi TRAV* /*TRBV* and *TRAC* /*TRBC* calls as positive alpha/beta evidence, and the new family-wide *TRGC* /*TRDC* result as a separate gamma/delta constant-region analysis. The family-wide Gobiesocidae matrix was therefore redrawn without the previous combined TCR alpha/beta/gamma/delta row, and no family-wide TCR absence was inferred. The gamma/delta result was discussed as a candidate component of the antibody-loss syndrome because Ig loss and B-cell-axis erosion now map to the Gobiesocidae root, but the TRG/TRD result was not elevated to an Ig-equivalent root loss because TRGC-like exons remain detectable in several genomes.

### 4.5. RAG2 PHD-domain motif comparison

RAG2 protein sequences were compared across the seven Gobiesocidae assemblies and eight blenniiform outgroups. For Gobiesocidae annotations lacking a direct RAG2 model, the region was recovered by protein-guided genomic search and translated manually. Candidate RAG2 open reading frames were checked for internal stop codons before motif comparison. The PHD-domain zinc-binding motif was aligned around the conserved GYWIKCC region, with residue numbering following *Lipophrys pholis* RAG2. Alignments were generated and inspected using standard multiple-sequence alignment workflows compatible with MAFFT-style protein alignment [26]. The resulting comparison was used as a protein-level marker associated with Ig-gene status, not as a stand-alone test of RAG2 function.

### 4.6. CD79A/CD79B synteny checks

Local synteny around *CD79A* and *CD79B* was inspected using the *S. fasciatus*-to-*G. willdenowi* homology table and the corresponding NCBI GFF annotations. For each locus, the *S. fasciatus* target gene and neighboring protein-coding genes were extracted from the chromosome-level annotation. Orthologous flanking genes in *G. willdenowi* were identified from reciprocal protein homology where available and then mapped back to the *G. willdenowi* GFF. The target gene was considered syntenically unsupported in *G. willdenowi* when the flanking genes were retained in the expected local context but no corresponding protein or genomic hit was detected for the *CD79* gene. The seven-species Gobiesocidae presence/absence matrix was displayed alongside the synteny panel as family-level context.

For the focused CD79 mechanism check, the *S. fasciatus* CD79A protein (XP_029958779.1) and CD79B protein (XP_029953778.1) were searched against the full *G. willdenowi* genome with tblastn using a permissive remnant-oriented configuration (word size 2, composition filtering disabled, and e-value reporting up to 1000). The same query set was searched separately against the extracted *Gouania* syntenic intervals spanning *LIPEB* -*ARHGEF1* for CD79A and *SCN4AA*-*PLEKHM1* for CD79B. Hits were considered candidate remnants only if they reached e-value ≤ 10^−3^ and query coverage ≥ 30%. The machine-readable query and result files are supplied as cd79_salarias_queries.fa, cd79_vs_gouania_genome.tblastn.tsv, cd79_vs_gouania_cd79a_interval.tblastn.tsv, cd79_vs_gouania_cd79b_interval.tblastn and cd79_mechanism_summary.tsv.

### 4.7. IgH expected-locus check

The expected IgH locus was evaluated by transferring the flanking-gene context from the annotated *S. fasciatus* IgH region on chromosome 4 (NC_043748.1). The positive-control *S. fasciatus* interval was defined from the *CABP5* -like gene at 15,905,267–15,907,701 to *NACC1A* at 16,910,026–16,921,168; this span contains annotated IgH constant-region genes, IgH-like coding genes, and heavy-chain V segments. The orthologous *G. willdenowi* flanking genes were identified on chromosome 8 (NC_041051.1), where *NACC1A* lies at 9,601,859–9,632,288 and the nearest *CABP5* -like annotation lies at 9,788,283–9,792,002. Gene annotations in this interval were inspected directly for IgH, IgM/IgD/IgT, heavy-chain V-segment, and heavy-chain constant-region terms. Fourteen annotated *S. fasciatus* IgH constant-region/heavy-chain proteins from the chromosome 4 IgH cluster were also used as tblastn queries against the extracted *Gouania* chromosome 8 interval and against the full *Gouania* genome. The result files are supplied with the igh_ prefix, together with gouania_igh_expected_interval.fa and igh_locus_mechanism_summary.tsv.

### 4.8. TAP and MHC class I review

TAP calls were checked directly in the *G. willdenowi* protein and GFF annotations, the refined immune homology table, and the *S. fasciatus*-*Gouania* gene-status comparison. Rows for *TAP1* and *TAP2A* were excluded from the displayed matrix because locus-level review contradicted the initial symbol-level absence calls. MHC class I annotations were inspected only as supporting context for functional interpretation; they were not used as proof of antigen presentation without expression or protein-level assays.

### 4.9. MHC class II pathway screen

To distinguish structural MHC class II retention from regulatory disruption, representative MHC class II and antigen-presentation proteins from annotated *Salarias*, *Blennius*, *Lipophrys*, and *Gouania* references were searched against the seven Gobiesocidae genomes with tblastn. Presence calls were based on clustered HSP support for MHC class II alpha and beta chains, *CD74*, *CTSS*, *CIITA*, RFX-family components, and NF-Y subunits. Weak or fragmentary hits were retained in the evidence table but were not encoded as intact presence calls.

### 4.10. Innate immune annotation survey

To test whether *G. willdenowi* retains immune systems that could provide candidate compensatory capacity, I screened the RefSeq feature table for protein-coding gene rows annotated as innate immune receptors, complement components, lectins, interferon-related genes, TRIM-family antiviral genes, and antimicrobial peptides. Counts were based on gene rows, not transcript or CDS isoforms. Product-name matches were used only as an annotation-level survey; they were not treated as orthology-validated expansions or functional proof of compensation. The same string-based counting scheme was also applied to *S. fasciatus* as a rough annotation reference.

### 4.11. Homology refinement of candidate gene absences

Candidate missing genes in *G. willdenowi* were refined by comparing candidate symbols against homologous sequences in related teleost references. Genes were classified as rescued by convergent homology, rescued by a single reference, weakly supported, ambiguous, or absent with high confidence. Immune genes were then reviewed separately to identify the subset of losses that remained robust after homology rescue.

### 4.12. Salarias-Gouania genomic comparison

Named protein-coding genes from *S. fasciatus* were compared to *G. willdenowi* using protein-level homology and genome-level alignment. Candidate genes lacking protein support and lying in regions with little detectable genomic alignment were classified as high-confidence missing-DNA candidates. Candidate blocks were defined by consecutive absent genes on *Salarias* chromosomes and classified as single-gene losses, small blocks, or large blocks.

### 4.13. Gobiesocidae target-gene matrix

A targeted immune-gene matrix was compiled across the seven Gobiesocidae species for 40 immune-associated genes selected from the *Salarias*-anchored missing-DNA comparison. Presence was assigned by tblastn only when the best hit met the predefined threshold (e-value ≤ 10^−5^, percent identity ≥ 25%, and query coverage ≥ 40%). This threshold was chosen for sensitivity and is permissive for vertebrate comparative genomics; it can include distant paralogs or shared-domain hits. For that reason, binary positives were treated as screening signals rather than definitive orthology calls, and broad conclusions were limited to repeated all-zero patterns or manually checked high-value loci. Weak hits below this threshold were retained in the evidence table but encoded as zero in the binary matrix. Genes with all-zero binary patterns were interpreted as root-compatible candidate absences; genes with one or more strong hits were treated as patchy, retained, or orthology-sensitive rather than forced into the root-loss set.

## 5. Acknowledgments

The author thanks the genome-sequencing groups that generated the chromosome-level Gobiesocidae assemblies used in this study. Computational work used publicly available NCBI GenBank and RefSeq data.

## Author contributions

F.G.D. designed the study, performed the manual immunoglobulin searches and comparative analyses, interpreted the data, and wrote the manuscript.

## Declaration of competing interest

The author declares no competing interests.

## Data and code availability

The manuscript uses publicly available genome assemblies and local analysis outputs listed in the Materials and Methods and supplementary tables. The LaTeX source, figure-generation script, figures, and machine-readable tables are provided in the manuscript directory.

## Supplementary figure legends

**Supplementary Fig. S1.**
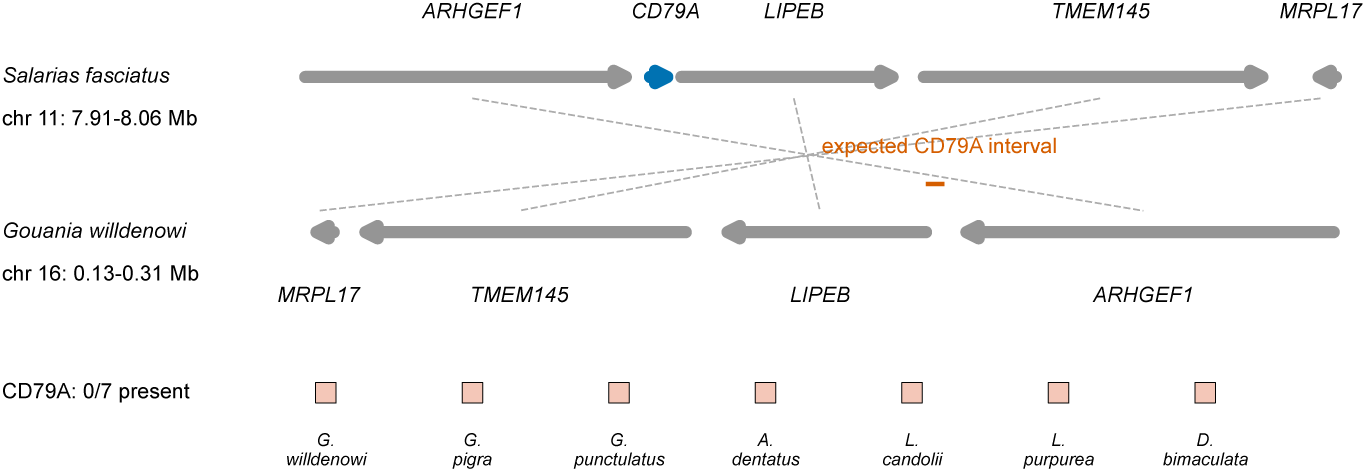
Synteny check for the *CD79A* locus. *S. fasciatus* retains *CD79A* between conserved flanking genes, including *ARHGEF1* and *LIPE*. In *G. willdenowi*, the orthologous flanks are retained on chromosome 16, but the expected *CD79A* interval has no detected protein or genomic hit. The lower matrix row shows the current Gobiesocidae target-gene matrix, in which *CD79A* is scored absent in all seven sampled species.

**Supplementary Fig. S2.**
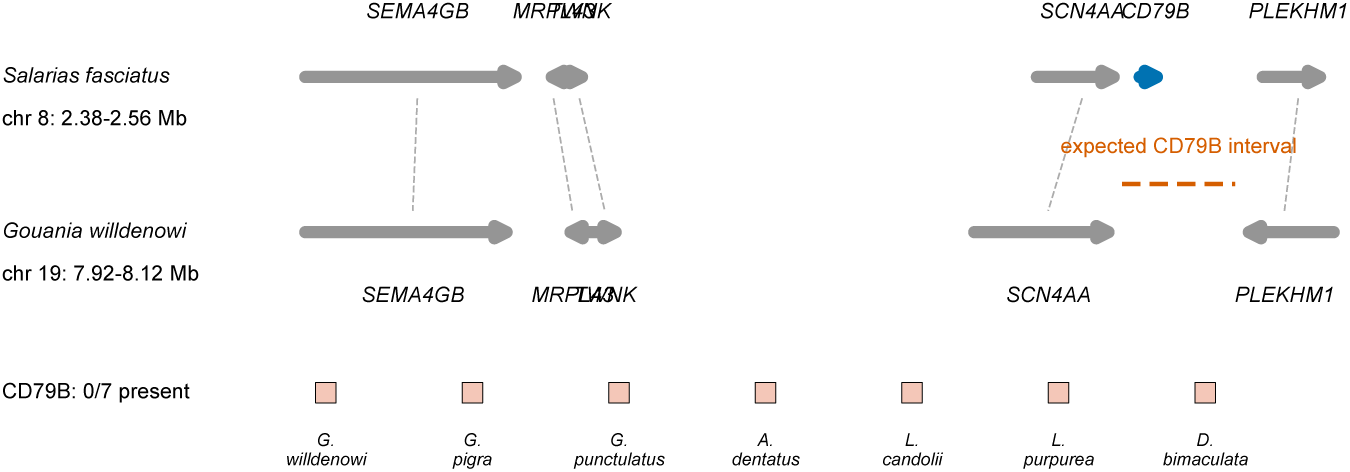
Synteny check for the *CD79B* locus. *S. fasciatus* retains *CD79B* in a conserved neighborhood including *SCN4A* and the downstream *PLEKHM1* side of the locus in the current *Salarias*-anchored comparison. In *G. willdenowi*, the orthologous flanks are retained on chromosome 19, but *CD79B* itself is not detected. The lower matrix row shows the current Gobiesocidae target-gene matrix, in which *CD79B* is scored absent in all seven sampled species.

**Supplementary Fig. S3.**
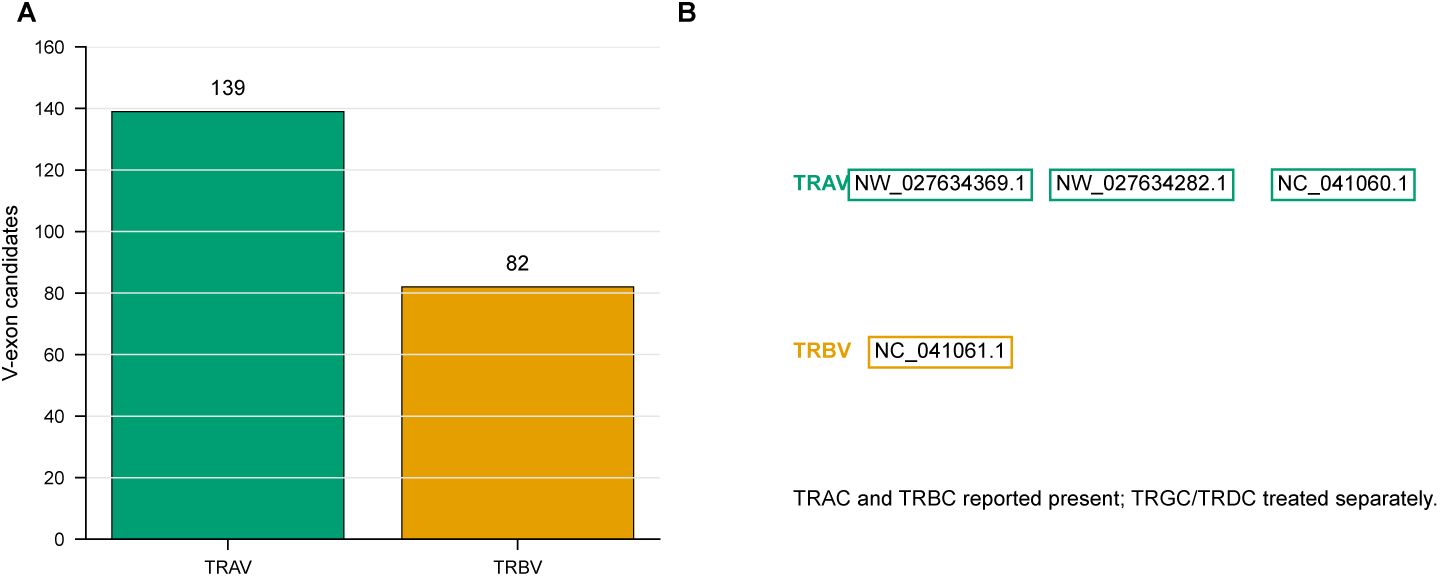
TCR alpha/beta evidence in *Gouania willdenowi*. Summary of prior *TRAV* /*TRBV* V-exon calls in *G. willdenowi*. The prior study reported 139 *TRAV* candidates on scaffold 33, scaffold 258, and Super Scaffold 5, and 82 *TRBV* candidates on Super Scaffold 7. These positive alpha/beta calls are kept separate from the family-wide *TRGC* /*TRDC* constant-region screen described in the main text.

## Supplementary tables

**Table 6:** TRG/TRD constant-region screen including the positive outgroup control. The table summarizes the *S. fasciatus* outgroup and all seven Gobiesocidae genomes. Accepted candidates are classifier-positive splice-bounded exons with probability at least 0.5. Strict direct tblastn support required e-value ≤ 10^−5^, identity ≥ 30%, query coverage ≥ 50%, and bit score ≥ 45. Best-hit values are shown as percent identity/query coverage/bit score. The full machine-readable table is supplied as table_s1_trg_trd_screen.tsv.

| Species | TRGC accepted | TRGC strict | TRGC hits | TRGC best | TRDC accepted | TRDC strict | TRDC hits | TRDC best |
| --- | --- | --- | --- | --- | --- | --- | --- | --- |
| *S. fasciatus* | 325 | 1 | 3270 | 100.0/100/211.0 | 0 | 0 | 27 | 74.1/33/43.9 |
| *G. willdenowi* | 0 | 0 | 32 | 32.2/94/43.5 | 0 | 0 | 2 | 45.7/44/40.0 |
| *G. pigra* | 0 | 0 | 14 | 32.3/97/42.0 | 0 | 0 | 2 | 45.7/44/40.0 |
| *G. punctulatus* | 0 | 0 | 21 | 31.0/92/41.6 | 0 | 0 | 0 | – |
| *A. dentatus* | 2 | 0 | 6 | 31.2/84/46.2 | 0 | 0 | 2 | 31.7/71/40.4 |
| *L. candolii* | 2 | 0 | 1 | 29.0/94/41.6 | 0 | 0 | 2 | 35.3/66/42.7 |
| *L. purpurea* | 1 | 0 | 14 | 30.3/84/41.2 | 0 | 0 | 4 | 35.1/75/38.9 |
| *D. bimaculata* | 2 | 0 | 1 | 28.6/87/40.4 | 0 | 0 | 0 | – |

**Table 7:** Scope of the immunoglobulin pseudogene/remnant interpretation. The table separates evidence for absence of canonical coding Ig loci from the unperformed, stricter question of whether all eroded molecular remnants have also disappeared. The machine-readable version is supplied as table_s2_ig_pseudogene_scope.tsv.

| Item | Evidence | Interpretation |
| --- | --- | --- |
| *S. fasciatus* IgH control | The NCBI feature table contains an IgH-rich region on chromosome 4, including immunoglobulin mu heavy-chain-like C-region genes and many Ig heavy-chain V-region genes around NC_043748.1:15.9–16.9 Mb. | Positive outgroup context for intact coding antibody loci. |
| *G. willdenowi* annotation check | Feature-table searches recover generic immunoglobulin-superfamily/domain genes and unrelated terms such as IGHM-enhancer binding proteins, but no annotated canonical IgH or IgL antigen-receptor locus. | Supports absence of annotated coding Ig loci, but does not alone exclude eroded pseudogene fragments. |
| Seven Gobiesocidae CH/V screens | CHfinder did not recover accepted IgM, IgD, or IgT heavy-chain constant exons; Vgeneextractor did not recover accepted light-chain V segments in the seven sampled Gobiesocidae genomes. | Supports family-wide absence of canonical coding IgH/IgL loci. |
| TRGC-like exons | TRGC-like subthreshold or accepted constant exons persist in several Gobiesocidae genomes while TRDC and TRGV remain unsupported. | Shows why antigen-receptor remnants must be discussed separately from intact functional loci. |
| Pseudogene scope | No systematic all-frame low-identity remnant screen with frameshift and stop-codon scoring was performed across all seven genomes. | The manuscript should state absence of canonical functional Ig loci, not complete absence of every possible molecular remnant. |

## Notes

### Competing Interest Statement

The authors have declared no competing interest.

## References

[1] M. F. Flajnik, A cold-blooded view of adaptive immunity, Nature Reviews Immunology 18 (7) (2018) 438–453. doi:10.1038/s41577-018-0003-9.

[2] M. F. Flajnik, M. Kasahara, Origin and evolution of the adaptive immune system: genetic events and selective pressures, Nature Reviews Genetics 11 (1) (2010) 47–59. doi:10.1038/nrg2703.

[3] M. D. Cooper, M. N. Alder, The evolution of adaptive immune systems, Cell 124 (4) (2006) 815–822. doi:10.1016/j.cell.2006.02.001.

[4] J. O. Sunyer, Fishing for mammalian paradigms in the teleost immune system, Nature Immunology 14 (4) (2013) 320–326. doi:10.1038/ni.2549.

[5] L. Tort, J. C. Balasch, S. Mackenzie, Fish immune system. a crossroads between innate and adaptive responses, Inmunología 22 (3) (2003) 277–286.

[6] D. Gómez, J. O. Sunyer, I. Salinas, The mucosal immune system of fish: the evolution of tolerating commensals while fighting pathogens, Fish & Shellfish Immunology 35 (6) (2013) 1729–1739. doi:10.1016/j.fsi.2013.09.032.

[7] S. Mirete-Bachiller, D. N. Olivieri, F. Gambón-Deza, Gouania willdenowi is a teleost fish without immunoglobulin genes, Molecular Immunology 132 (2021) 102–107. doi:10.1016/j.molimm.2021.01.022.

[8] J. C. Briggs, A monograph of the clingfishes (order Xenopterygii), Stanford Ichthyological Bulletin 6 (1955) 1–224.

[9] K. W. Conway, D. Kim, L. Rüber, H. S. Espinosa Pérez, P. A. Hastings, Molecular systematics of the New World clingfish genus *Gobiesox* (Teleostei: Gobiesocidae) and the origin of a freshwater clade, Molecular Phylogenetics and Evolution 112 (2017) 138–147. doi: 10.1016/j.ympev.2017.04.024.

[10] K. W. Conway, B. B. Collette, P. A. Hastings, Molecular phylogenetics of the clingfishes (Teleostei: Gobiesocidae)—Implications for classification, Copeia 108 (4) (2020) 827–843. doi:10.1643/ci2020054.

[11] P. Wagner, M. Kovačić, C. Fruciano, O. Seehausen, J. Pfaender, Diversification in gravel beaches: A radiation of interstitial clingfish (*Gouania*, Gobiesocidae) in the Mediterranean Sea, Molecular Phylogenetics and Evolution 139 (2019) 106525. doi:10.1016/j.ympev.2019.106525.

[12] P. Wagner, M. Kovačić, O. Seehausen, J. Pfaender, Unravelling the taxonomy of an interstitial fish radiation: Three new species of *Gouania* (Teleostei: Gobiesocidae) from the Mediterranean Sea and redescriptions of *G. willdenowi* and *G. pigra*, Journal of Fish Biology 97 (3) (2020) 694–726. doi:10.1111/jfb.14558.

[13] T. J. Near, R. I. Eytan, A. Dornburg, K. L. Kuhn, J. A. Moore, M. P. Davis, P. C. Wainwright, M. Friedman, W. L. Smith, Resolution of ray-finned fish phylogeny and timing of diversification, Proceedings of the National Academy of Sciences 109 (34) (2012) 13698–13703. doi: 10.1073/pnas.1206625109.

[14] H.-C. Lin, P. A. Hastings, Phylogeny and biogeography of a shallow water fish clade (Teleostei: Blenniiformes), BMC Evolutionary Biology 13 (2013) 210. doi:10.1186/1471-2148-13-210.

[15] W. Reith, S. LeibundGut-Landmann, J.-M. Waldburger, Regulation of MHC class II gene expression by the class II transactivator, Nature Reviews Immunology 5 (10) (2005) 793–806. doi:10.1038/nri1708.

[16] A. G. W. Matthews, A. J. Kuo, S. Ramón-Maiques, S. Han, K. S. Champagne, D. Ivanov, M. Gallardo, D. Carney, P. Cheung, D. N. Ciccone, K. L. Walter, P. J. Utz, Y. Shi, T. G. Kutateladze, W. Yang, O. Gozani, M. A. Oettinger, RAG2 PHD finger couples histone H3 lysine 4 trimethylation with V(D)J recombination, Nature 450 (7172) (2007) 1106–1110. doi:10.1038/nature06431.

[17] S. Ramón-Maiques, A. J. Kuo, D. Carney, A. G. W. Matthews, M. A. Oettinger, O. Gozani, W. Yang, The plant homeodomain finger of RAG2 recognizes histone H3 methylated at both lysine-4 and arginine-2, Proceedings of the National Academy of Sciences 104 (48) (2007) 18993–18998. doi:10.1073/pnas.0709170104.

[18] N. Shimazaki, A. G. Tsai, M. R. Lieber, H3K4me3 stimulates the V(D)J RAG complex for both nicking and hairpinning in trans in addition to tethering in cis: implications for translocations, Molecular Cell 34 (5) (2009) 535–544. doi:10.1016/j.molcel.2009.05.011.

[19] M. Nei, A. P. Rooney, Concerted and birth-and-death evolution of multigene families, Annual Review of Genetics 39 (2005) 121–152. doi:10.1146/annurev.genet.39.073003.112240.

[20] M. Reth, Antigen receptors on B lymphocytes, Annual Review of Immunology 10 (1992) 97–121. doi:10.1146/annurev.iy.10.040192.000525.

[21] T. Kurosaki, H. Shinohara, Y. Baba, B cell signaling and fate decision, Annual Review of Immunology 28 (2010) 21–55. doi:10.1146/annurev.immunol.021908.132541.

[22] F. Mackay, J. L. Browning, BAFF: a fundamental survival factor for B cells, Nature Reviews Immunology 2 (7) (2002) 465–475. doi:10.1038/nri844.

[23] J. Neefjes, M. L. M. Jongsma, P. Paul, O. Bakke, Towards a systems understanding of MHC class I and MHC class II antigen presentation, Nature Reviews Immunology 11 (12) (2011) 823–836. doi:10.1038/nri3084.

[24] J. S. Blum, P. A. Wearsch, P. Cresswell, Pathways of antigen processing, Annual Review of Immunology 31 (2013) 443–473. doi:10.1146/annurev-immunol-032712-095910.

[25] S. F. Altschul, T. L. Madden, A. A. Schaffer, J. Zhang, Z. Zhang, W. Miller, D. J. Lipman, Gapped BLAST and PSI-BLAST: a new generation of protein database search programs, Nucleic Acids Research 25 (17) (1997) 3389–3402. doi:10.1093/nar/25.17.3389.

[26] K. Katoh, D. M. Standley, MAFFT multiple sequence alignment software version 7: improvements in performance and usability, Molecular Biology and Evolution 30 (4) (2013) 772–780. doi:10.1093/molbev/mst010.

